# SCIP: a self-paced, community-based summer coding program creates community and increases coding confidence, lessons learned from the pandemic

**DOI:** 10.1101/2022.12.27.521952

**Authors:** Rochelle-Jan Reyes, Olivia Pham, Ryan Fergusson, Niquo Ceberio, Candace Clark, C Sarah Cohen, Megumi Fuse, Pleuni Pennings

## Abstract

In 2020, many students lost summer opportunities due to the COVID-19 pandemic. We wanted to offer students an opportunity to learn computational skills and be part of a community while they were stuck at home. Because the pandemic was very isolating, it was important to support students to learn and build community online. We used lessons learned from literature and our own experience to design, run and test an online program for students called the Science Coding Immersion Program (SCIP). In our program, students worked in small teams for 8 hours a week spread over the week, with one participant as the team leader and Zoom host. Teams worked on an online R or Python class at their own pace with support on Slack from the organizing team. For motivation and career advice, we hosted a weekly webinar with guest speakers. We used pre- and post-program surveys to determine how different aspects of the program impacted students. We were able to recruit a large and diverse group of participants who were happy with the program, found community in their team, and improved their coding confidence. We hope that our work will inspire others to start their own version of SCIP.

## INTRODUCTION

In the summer of 2020, the COVID-19 pandemic caused many summer research programs and internships to be delayed or canceled. A survey in the United States reported that 44.5% of research opportunities for undergraduate students were canceled (Grineski, 2020), which left many students without the opportunities for learning and career advancement. To combat this loss of opportunities, some undergraduate research institutions adjusted their summer programs to become virtual or created new online programs (Noriega 2020, Valerie Sloan 2020, Afghani 2021, Yowler et al. 2021).

At San Francisco State University (SF State) in-person learning was halted in March of 2020, students were no longer allowed to work in labs. It became clear that campus would not be open during the summer. The campus closure led to a huge loss of opportunity for our students. To provide our students a way to learn new skills despite the campus closure, we created a new virtual coding program, partly based on our previous experience with an on-campus summer coding program for non-computer science undergraduates (Pennings et al., 2020). With the new, online program, we wanted to provide a learning and leadership experience for students who had lost STEM opportunities due to the campus closure. The learning objective for this program was to increase coding confidence and enthusiasm amongst the participants. This broad learning outcome meant that multiple online coding courses could be used within our setting. In addition, we aspired to provide a sense of community for the participants, particularly since so many students had lost their university community when they were forced to move away from campus. Thus, the Science Coding Immersion Program (SCIP) was established, a community-based coding program designed for undergraduate and master’s students in Biology, Chemistry and Biochemistry. We had three major aims for SCIP: (1) to create a virtual workspace structure that could work while students were sheltered in place during the pandemic, (2) to provide a community for the students, and (3) to improve student confidence in coding as a scientific skill (Figure 1A).

**Figure 1.**
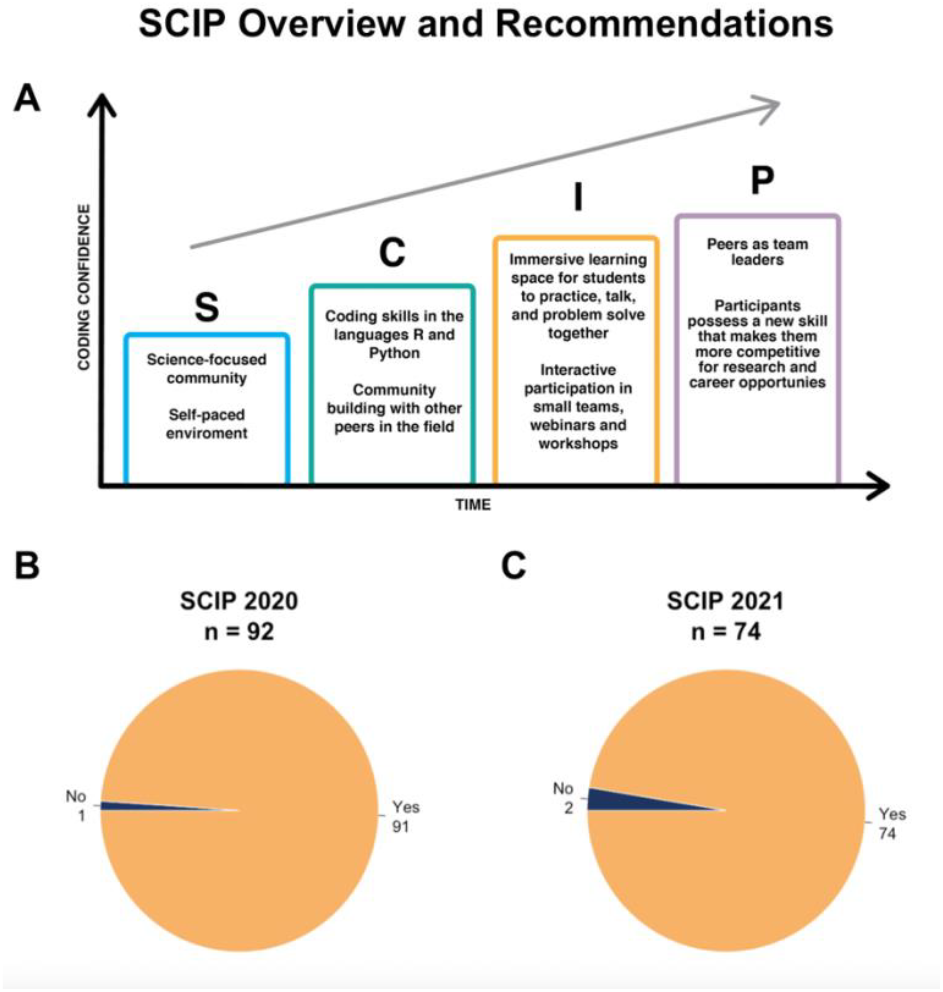
(A) Overall plan for design of the Science Coding Immersion Program (SCIP). (B) In the post-program assessment, SCIP participants from Summers 2020 and 2021 were asked whether they would recommend SCIP to others. In 2020, 91 out of 92 who answered the question responded that “Yes”, they would recommend SCIP, while in 2021, 74 out of 76 answered “Yes”.

This paper will explicitly describe how we developed a self-paced and community-based immersive virtual coding environment (Figure 1A), and how we assessed satisfaction and confidence of the participants. Overall, most participants polled were enthusiastic about their experience and would recommend this program to colleagues (Figure 1B). Given the student responses and the demographics of the student body, we believe that building these programs will support and speed up diversification of the sciences and will add to the research experiences students are receiving in the lab.

### DEVELOPING SCIP

SCIP had three major aims: (1) to create a virtual workspace structure that could work while students were sheltered in place during the pandemic, (2) to provide a scientific community for the students, and (3) to improve student confidence in coding as a scientific skill.

#### Aim 1: Create a virtual workspace structure

There were three main considerations in developing the logistics of the online SCIP program. 1) We wanted to offer flexible scheduling to accommodate other summer activities of participants, 2) we wanted to offer participants the choice to learn either Python or R, with the potential to include other courses in the future, and 3) we wanted to set up a system for successful communication between participants, team leaders and coordinators.

### Overall structure

The basic structure of SCIP was adapted from the ten-step guide to support non-computer science undergraduates in learning how to code in a summer program (Pennings et al., 2020). We then modified the structure to accommodate a larger number of participants with different schedules, last-minute planning for the organizing team and the participants, and a choice of different coding languages. Because the program was 100% virtual, we had to make sure we had a strategy for communication between participants, team leaders and coordinators. We also included a weekly webinar to showcase possible career paths for scientists with coding skills. Based on these ideas, we offered a program focused on Zoom meetings, staggered start dates (in 2020, but not 2021), two programming languages (a pilot study for ImageJ was offered in 2021, see Scheduling and Coding Languages), communication primarily on Slack, as well as extra readings, webinars and introductory datasets.

### Online Communication Platforms

To streamline communication, two communication platforms - *Slack* and *Zoom* - were chosen and used throughout the program (Figure 2). These platforms were chosen because they were free to use and most of the participants had prior experience with the use of these platforms at SF State. For example, SF State adopted Zoom conferencing services (Zoom Video Communications Inc., 2016) for virtual classes during the pandemic, and Slack (Slack Technologies) was integrated into some research labs and used by the majority of students funded by public (NIH and NSF) and private (Genentech Foundation) fellowships through the Student Enrichment Opportunities Office (https://seo.sfsu.edu/), as well as by the Department of Biology.

**Figure 2.**
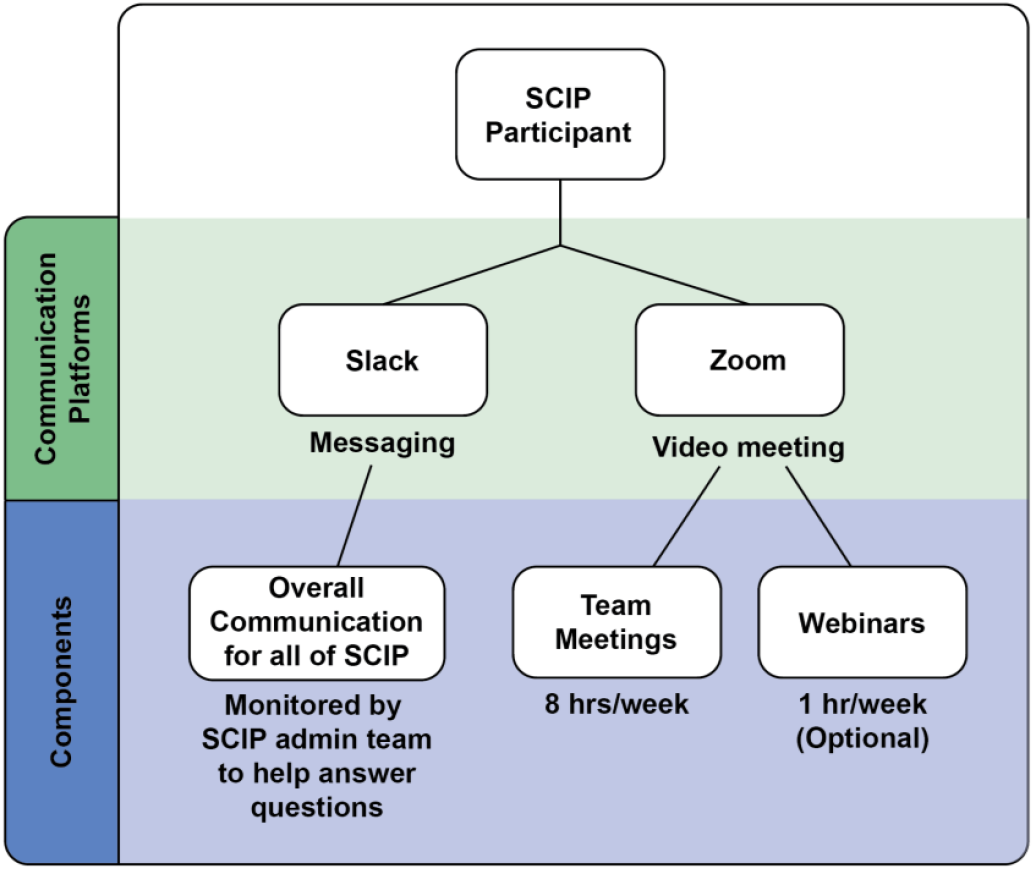
Communication platforms adopted in the program.

We used Slack as our primary communication tool between the organizing team and the participants. To assure efficient communication, all participants were required to join our Slack workspace before starting the program. The platform provided a space for the participants to communicate within their teams, with SCIP coordinators, and with any other participant in the program. General channels were accessible to everyone for webinar reminders and general announcements, and a question forum was available for participants to ask questions and share tips. In addition, a separate channel for each team was created in order for them to discuss coding and meeting topics outside of their team meetings.

SCIP participants and coordinators used Zoom for their team meetings and webinars. Teams used Zoom to meet virtually with their group and work on their coding courses four times a week. For each team, the team leader hosted the team meetings. In addition, Zoom webinars were held weekly to provide participants insight on research and career options within the Biology and Chemistry/Biochemistry fields that utilize coding. Guest speakers shared their experiences (and struggles) using coding skills in their job or graduate program, and speakers ranged from professors to SF State alumni who went on to graduate programs, research, and/or biotechnology jobs. The webinar topics were geared toward providing participants different perspectives on coding and its use in the sciences, to motivate them to learn and nurture their new skill and to build greater confidence. Thus, we used Zoom for team meetings and webinars, a Slack space for announcements and questions, and Google documents for instructions, all of which could be modified based on virtual platforms at any university.

### Scheduling and Coding Languages

SCIP was designed to be flexible with participants’ summer schedules and interests. Potential conflicts included summer courses, other virtual summer programs, part-time jobs and family commitments. When we had to introduce the course rapidly, as was done in 2020, we staggered the start dates, which allowed us to start off with a small number of participants as we were learning how to run the program. It also allowed the large volume of registrants to join at a time that best suited their schedules, and it ensured that participants who heard about the program late were still able to participate. With sufficient time to advertise and organize as such in 2021, we were able to begin all groups in the same week, with the same duration for the program. As will be discussed later, both were equally successful.

In 2020, participants chose their preferred Monday start date from a 5-week selection. The duration of the program varied from 8 weeks for the early start dates to 6 weeks for the later start dates. In 2021, all participants started on the first Monday of June in 2021 for 6 weeks of the program in total. Participants also indicated the times they had available (morning, afternoon, evening), the programming language they wanted to learn and indicated their experience level.

Each participant was also asked to indicate whether they were willing to be a team leader, which did not depend on any coding knowledge. It was communicated to the participants that team leaders were expected to have leadership experience (from a job, campus, church, etc.), but they didn’t need coding experience. They were asked to briefly discuss their previous leadership experience in a job, church, dorm etc. Most of our team leaders had little-to-no coding experience (survey data not shown). Zoom meetings were hosted and facilitated by these peer team leaders. Team leaders were learning the same material as everyone else in the team. Studies have shown that peer-led teams have a positive impact on participants’ performance, retention, and attitude (Batten & Ross, 2021; Quitadamo et al., 2009; Tien et al., 2002; Trujillo et al., 2015). To make sure the team leaders could be successful and didn’t need to spend extra time preparing for meetings, we created a schedule for them to use in the meetings with very detailed information for the first few days and less detail later on (Supplemental Document S1). Our material for team leaders included sample emails for them to send to their team, and checklists for what they were expected to do each week (Table 1).

**Table 1.**
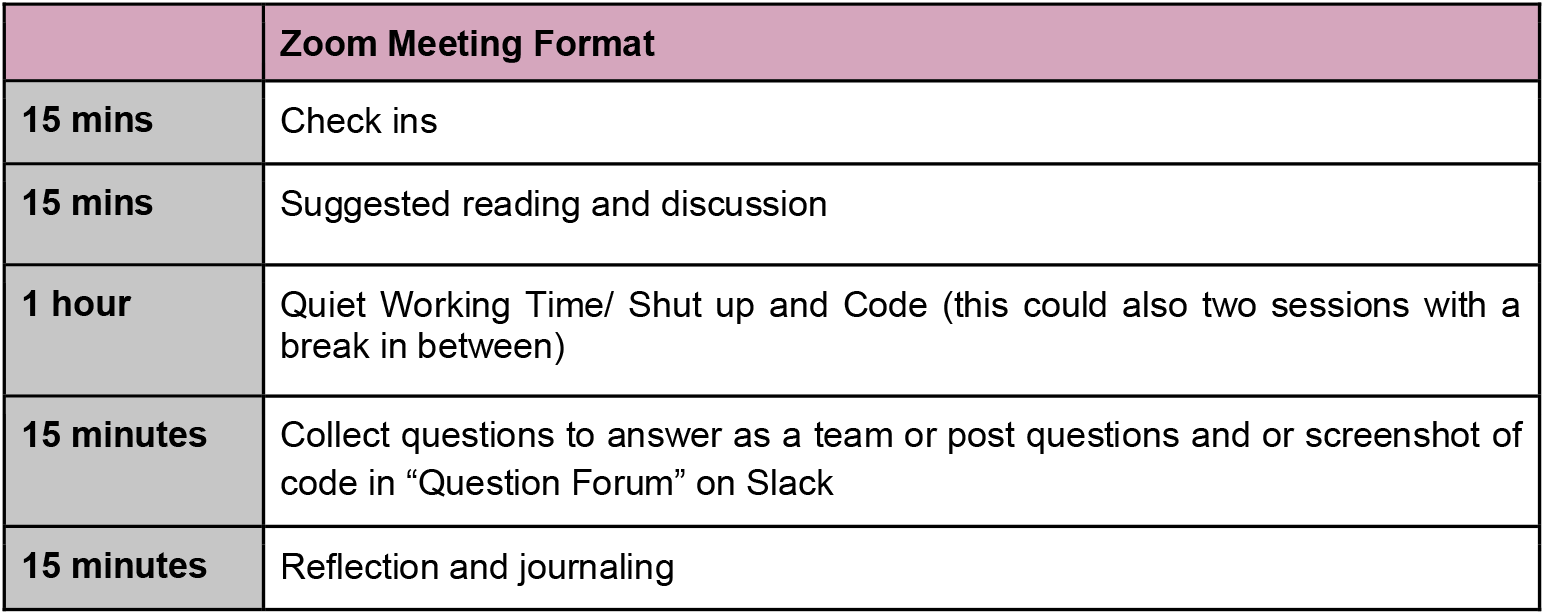
Suggested Zoom meeting format. This table outlines a suggested format that teams could follow during their meeting time. Each team had 4 such meetings each week.

We created teams based on programming language preference, time availability, and experience level. Participants were grouped in teams of 4-8 participants with a participant as team leader. Each team worked independently on a course that was chosen by us (Table 2). The courses we offered in 2020 were the free beginner Udacity courses of Data Analysis with R (Udacity: Data Analysis with R, n.d.) and Introduction to Python Programming (Udacity: Introduction to Python, n.d.). In addition to R and Python, we added an image processing option in 2021, using ImageJ as a learning platform (edX Image Processing and Analysis, n.d.).

**Table 2.**
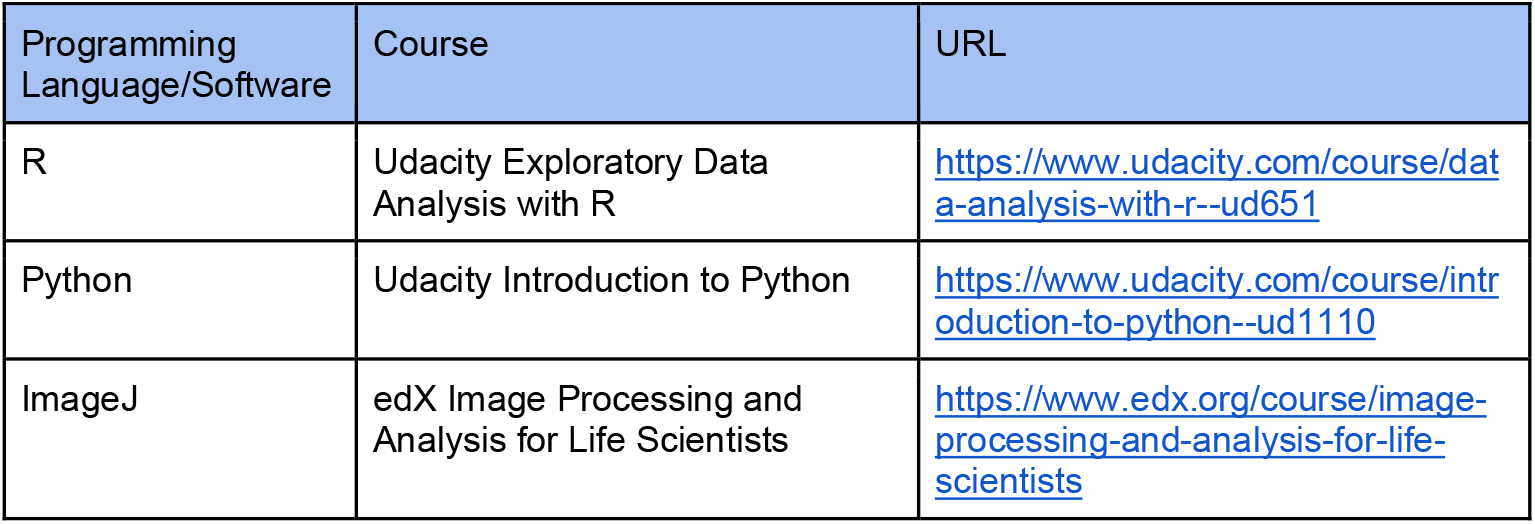
List of programming languages or software we offered as well as the source of these classes.

### Program Learning Structure

What sets SCIP apart from other online learning structures is the immersion aspect of the program, which allows participants to learn both as part of a community and individually. In the SCIP program, teams meet for 4 days a week for 2 hours per meeting (8 hours a week total) with about 4 of those hours designated as “quiet working time”. Participants are encouraged to work on their online course only during the “quiet working time” in their team meetings, and not outside of the team meetings. Therefore, participants learned new skills while they were in Zoom meetings with each other during the reserved meeting time, rather than working on it in their own time. We structured our program this way so that participants would never struggle with coding tasks on their own. The idea of having “quiet working time” was based on previous work in “Rules for a Bioinformatics Journal Club’’ (Lonsdale et al., 2016) and “Ten simple rules for an inclusive summer coding program for non-computer-science undergraduates’’ (Pennings et al., 2020).

Team leaders were instructed to make sure that each Zoom meeting consisted of time to talk (check-ins, problem solving, brainstorming etc.) and time to study (read, view course videos, do coding exercises) while everyone in the Zoom meeting was muted (Table 1). Teams were not restricted to the suggested format and were free to create their own schedule as long as the following topics were covered every Zoom meeting: (1) brief check-ins to share how they were doing / how they felt about coding that day, (2) weekly suggested reading and discussion, (3) a total of 1 hour of quiet working time or “Shut up and Code” time per session, (4) collection of questions to answer as a team or post on Slack and (5) reflections.

A core value of SCIP within this group setting was that participants were encouraged to learn how to code in a self-paced manner without deadlines or tests. Self-paced learning was encouraged for each individual student, though some teams tried to all work at the same pace. During the Zoom meetings, participants were able to ask each other for help regardless of where they were in the course. Because of Slack, they were also able to talk with peers, team leaders and SCIP coordinators. This allowed participants who were ahead in the course to provide support for those who were in earlier parts of the lessons.

### Assessing effectiveness of Zoom and Slack communicating

Evaluation of the effectiveness of various aspects of the program was analyzed through data from two participant surveys: a pre- and a post-program survey. Participation in the survey was voluntary, and all participant survey responses were collected anonymously. Because group start dates were staggered in 2020, the pre-assessment survey was given after the last offered start date of the program and the post-assessment survey was given at the end of the entire program. For 2021, pre-assessment was given on the first day of the program since all participants started on the same day and post-assessment was sent out at the end of the program.

Pre- and post-assessment surveys were administered via Google forms, to obtain evaluation of the program as well as its components, through the perspective of the participants (See Supplemental Document S2 and S3). In particular, we evaluated whether the communication platforms Zoom and Slack were effective in our program (Aim 1), whether the demographic and participation data reflected that of the biology and chemistry/biochemistry departments (see Aim 2) and whether the participants’ overall confidence in coding changed by participating in SCIP (see Aim 3). Rating scale questions, in the form of Likert-type data, were asked in both assessments to evaluate confidence in coding as well as effectiveness of the program design. In the post-assessment, some questions were *open-ended* to elicit broader comments or suggestions about the SCIP program. The study was IRB exempt. Data and code for data analysis are available in a GitHub repository (Reyes & Pham, 2022).

Post-survey responses to whether the communication platforms Zoom and Slack were effective in our program, showed that 95.7% (88/92) of our participants felt that using the Zoom meetings with their teams were “Effective”, “Somewhat to Very Effective” or “Very Effective” in 2020, and 89.5% in 2021 (Figure 3A). For Slack, 91.3% (84/92) of the participants felt that the Slack workspace was “Effective” to “Very Effective” in their participation in SCIP in 2020 and 89.5% in 2021 (Figure 3B). The webinars likewise were considered “Effective” to “Very Effective” by 87.0% in 2020 and 93.4% in 2021 (Figure 3C). As a reminder, webinars were not mandatory activities.

**Figure 3.**
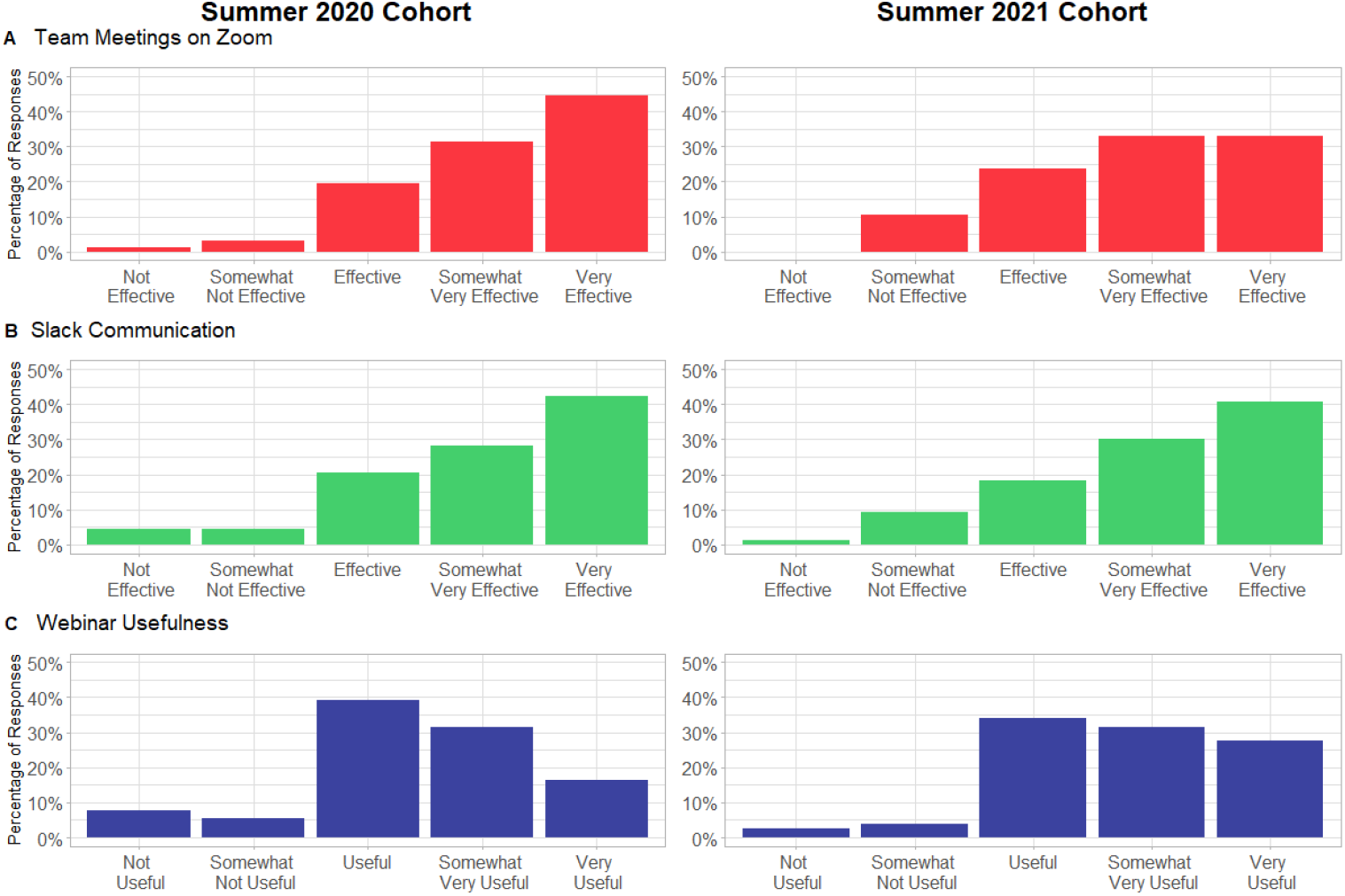
Survey-reported effectiveness of the use of (A) Zoom for team meetings, and (B) Slack workspace and (C) webinars, as media to communicate with team members as well as other SCIP participants and coordinators.

#### Aim 2: Providing a scientific community for the students

One goal of SCIP was to help students feel part of a community during a period when they were not on campus. Although labs began to open by 2021, in person classes at SF State were still very limited and a significant amount of time was spent with remote learning. We gauged the scientific community by analyzing (1) who joined SCIP, and (2) what participation and retention were in the SCIP program.

### SCIP participant pool mirrored SF State Biology demographics

In 2020, just over half (54.5%) of the participants were undergraduate students, 35.7% were master’s students and the remaining participants comprised of post-baccalaureates, faculty, and staff. In SCIP Summer 2021, undergraduate and master’s student counts were similar at 54.7% and 27.3%, respectively. Most participants were Biology and Chemistry/Biochemistry students (see Supplemental Figure S1).

Based on the pre-assessment survey, participants’ self-identified race and ethnicity mirrored SF State enrollment statistics in the Biology and Chemistry/Biochemistry departments for Spring 2020 (Office of Institutional Research, SF State: https://ir.sfsu.edu/; Figure 4). For both cohorts, the majority of participants were of Hispanic/Latinx (28.1 and 34.8% for 2020 and 2021 respectively) and Asian (22.3 and 35.2%) descent, followed by White and mixed-race. In comparison, of the approximately 800 students enrolled in SF State Biology (∼700) and Chemistry/Biochemistry (∼100) departments for spring 2020 data, 40.8% identified as Hispanic or Latinx descent. This somewhat lower number of Latinx students in our program compared to the university is in part because we allowed participants to check multiple ethnicities, in which case we counted them as mixed-race. Although the Black student population is not large in general at SF State, our demographics mirrored those in the Departments of Biology and Chemistry/Biochemistry and increased in 2021.

**Figure 4.**
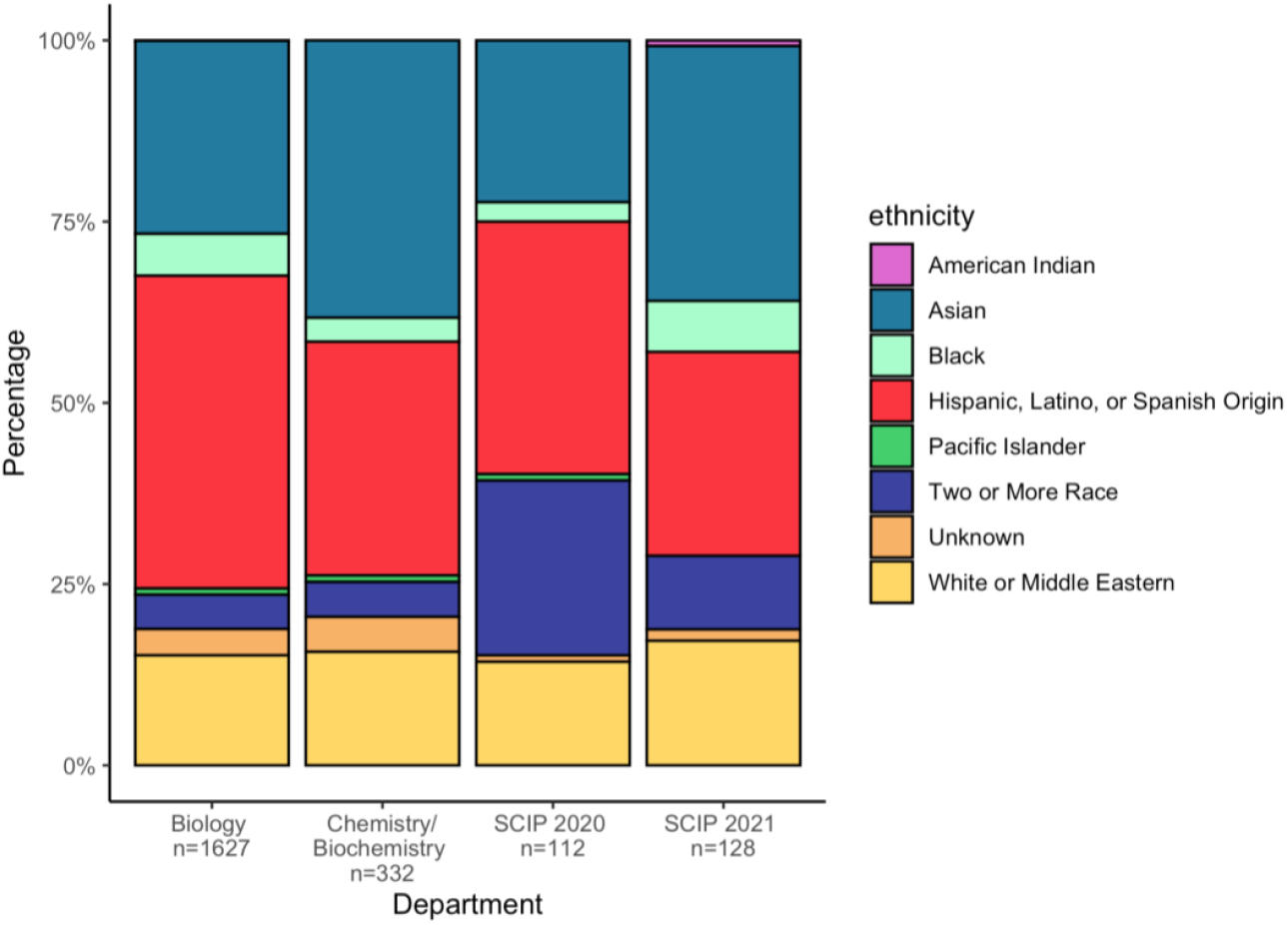
Race and Ethnicity of participants in SCIP mirrored the demographics of the Biology and Chemistry/Biochemistry departments. Participants who identified with more than one race and/or ethnicity were counted within the “Two or More Race” category. Spring 2020 data for Biology and Chemistry/Biochemistry were taken from SF State Institutional Research (www.ir.sfsu.edu).

### Participation and retention rates were strong

Participant information was based on the pre-assessment survey that was given prior to the completion of the program. There were 147 participants in SCIP 2020 and 138 in SCIP 2021 at the start of each program, as determined by the scheduling of all team members. Survey participation was optional and anonymous. Participation and retention rates were gathered based on personal communication with participants, survey responses, and presence of members in team photos (Table 3).

**Table 3.**
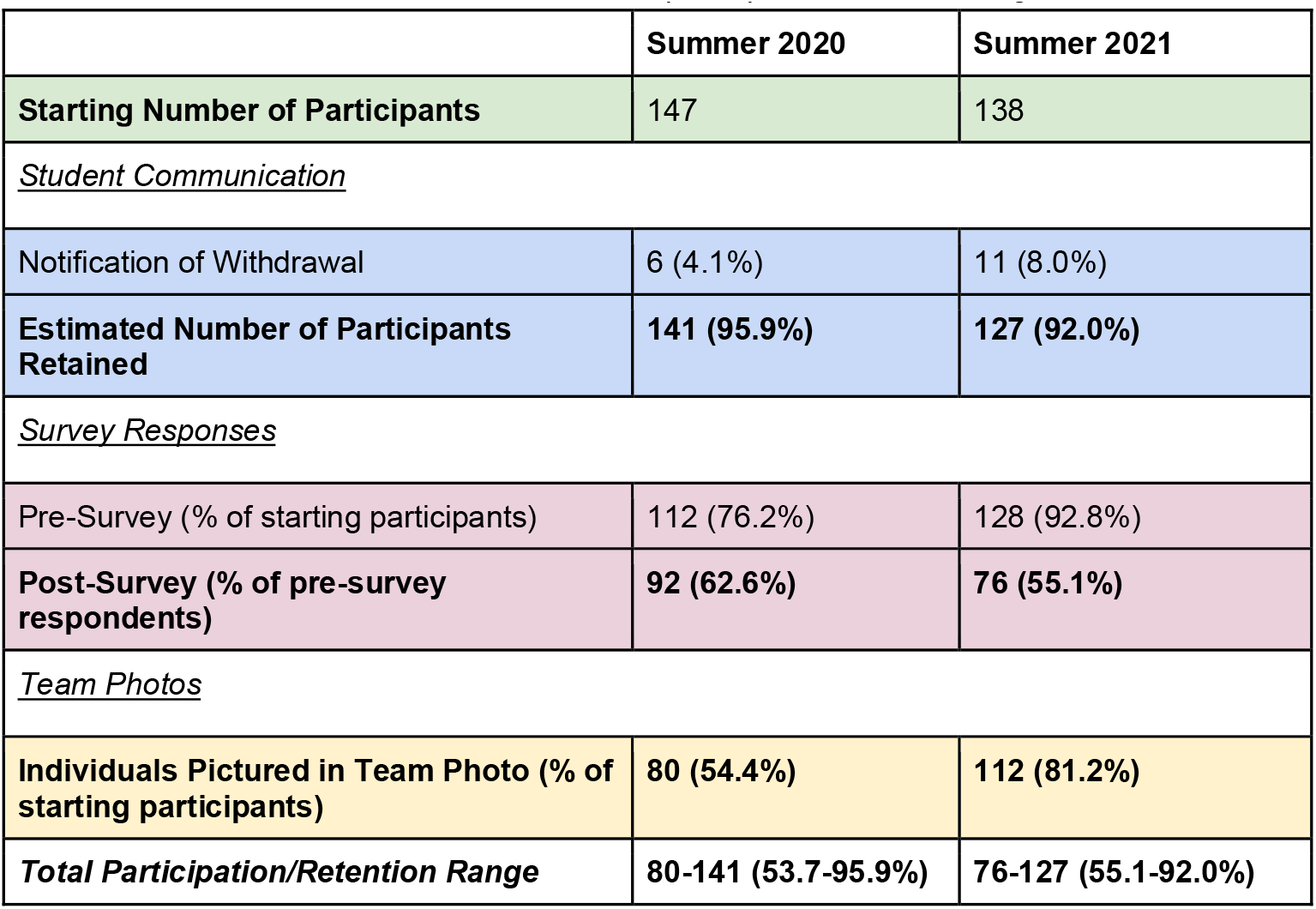
Participant Data and Estimates of Participant Retention from Student Communication regarding withdrawal from the program, Survey Responses, and presence in Team Photos. Bolded are the data utilized to calculate the total participation/retention range.

Based on communication with participants, 4.1% (6) of participants left the Summer 2020 cohort and 8.0% (11) left in Summer 2021. For the latter cohort, more information was available: 6 participants had scheduling conflicts and 5 had too large a summer workload.

The survey completion rate for the pre-assessment was 76.2% and 92.8% for 2020 and 2021, respectively (112 out of 147 total SCIP participants starting in 2020 and 128 out of 138 in 2021). The higher rate for 2021 was likely because in that year we specifically asked participants to do the survey on the first day of the program before they did anything else. Survey completion rate for the post-assessment survey was 62.6% and 55.1% for 2020 and 2021 respectively (92 out of 147 and 76 out of 138 of starting participants). These numbers show that in 2020, at least 62.6% of the participants stayed in the program until the end, while in 2021 at least 55.1% of the participants stayed in the program until the end. We believe that both of these numbers are underestimates of retention, because not all participants who stayed in the program filled out the post-assessment survey, especially in 2021.

Because we are aware that our survey responses may not accurately reflect participation rates, we also studied team photos to estimate participation rates. Team photos were taken in the last week and a half of the program and used to celebrate team success in working together throughout the summer. We assumed that the presence of a participant in a photo indicated that they were still active towards the end of the program. Referencing these photos, we saw that 80 participants were present on a team photo (53.7% of the 147 participants) in 2020. We also know that 7 teams with 40 participants did not submit team photos. Therefore, the 80 participants in the photos represent 74.8% of the 107 participants who could have been on a team photo. In 2021, we received photos from all teams and 81.2% (112 from the starting population) of participants were present in their team photo. We believe that this number (81.2%) is a reasonable estimate for retention for the 2021 program.

Based on these three metrics of student participation, we estimate that retention was strong, in the range of 53.7-95.9% for the Summer 2020 cohort and in the range of 55.1-92.0% for the Summer 2021 cohort. While these ranges are broad and there is substantial uncertainty about the retention in our program, we believe that the photos of all teams in 2021 give a good estimate of just over 80% retention at the end of the 6-week program.

### Participants felt a sense of community by the end of the program

To get a deeper understanding of what part of our program’s learning structure worked well for participants, we offered survey respondents an open-ended prompt to discuss what they enjoyed about the program. The wording of the prompt was as follows: “What did you like about SCIP?”. Survey participants’ responses were split into a list of words, and the frequency of the words was determined based on the number of times stated. Common conversational filler words such as “the”, “a”, and “is” were omitted. We created a word cloud to illustrate the most utilized words in the open-ended prompt (Figures 5A, C). We identified the top 15 most frequently stated words based on survey responses. Five of the top 15 frequently stated words associated with positive aspects of the program were “team”, “people”, “community”, “group”, and “collaboration”, with “team” being the most frequently used word in 2020 (Figure. 5A and B). In 2021, positive aspects of the program continued to involve similar words to those in 2020, such as “team” and “community” (Figure 5C, D). Many of the most common words in 2020 and 2021 show that the team-based aspect of the program was appreciated by the participants.

**Figure 5.**
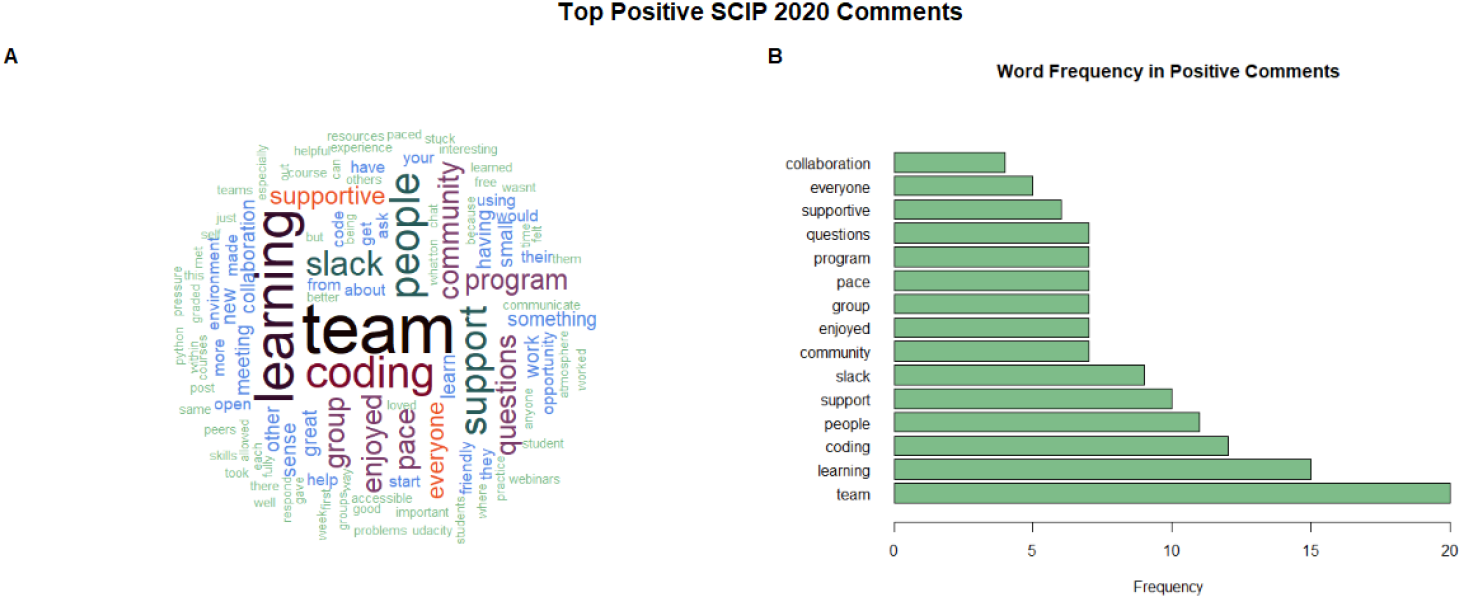
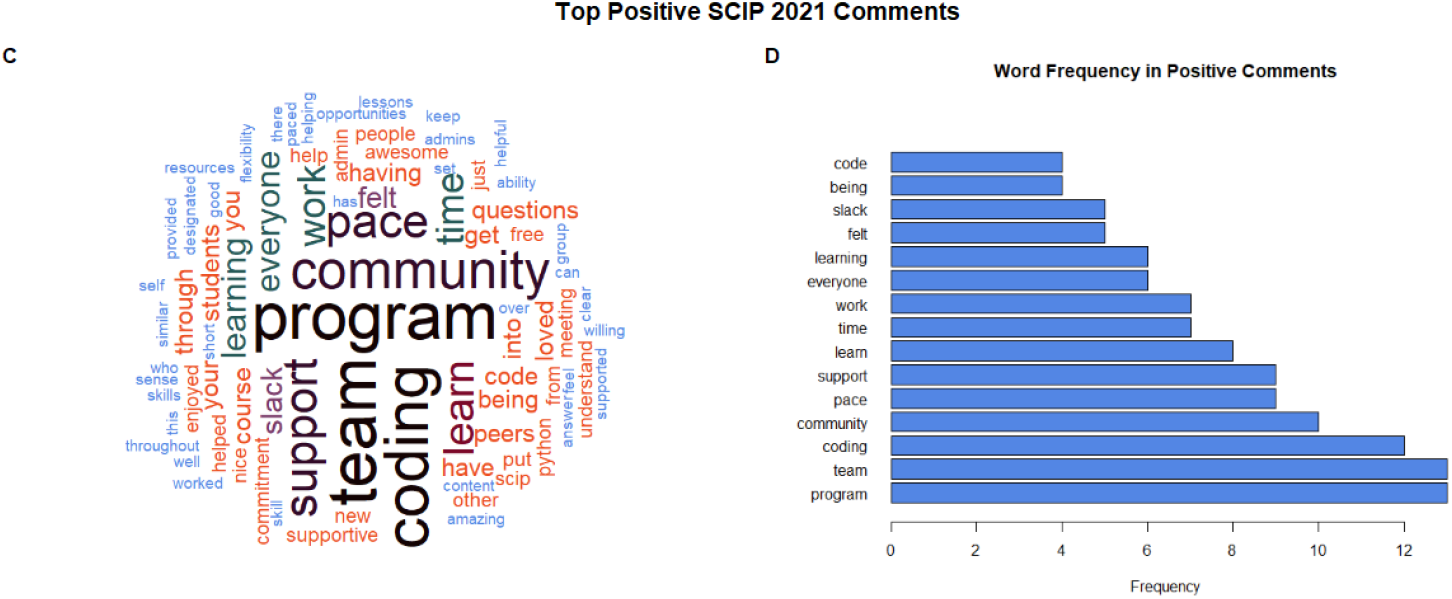
Words mentioned by participants in the post-assessment survey with the prompt, “What did you like about SCIP?”. Words were taken from the open-ended prompt and consolidated into a count of the words used, from highest count to lowest count. (A, C) Word Cloud of word frequency of the question with the most frequently used words in larger font size and location, in 2020 and 2021, respectively. (B, D) Bar plot of the frequency of words used in the question, in 2020 and 2021, respectively.

#### Aim 3: Improve student confidence in coding as a scientific skill

The third aim of SCIP was to provide participants an opportunity to learn a new skill to advance their careers or their research projects that involved coding-based tools. With this third aim, we hoped to increase the confidence of participants who were beginners in coding, while addressing both the lack of summer research opportunities as well as the possible shift in their research projects towards coding-based research approaches. This is especially useful because research fields like genetics, microbiology, neurobiology, ecology, and evolutionary biology often depend on programming to analyze big data (Stres & Kronegger, 2019). In addition, there are many opportunities in the computational biology and bioinformatics disciplines which integrate programming and statistical analyses with the life sciences (Ouzounis & Valencia, 2003). With the development of these disciplines and the need for students to learn the skills necessary to enter the field, our intent was to provide undergraduate and graduate students an opportunity to learn programming skills for their research and careers. Although we initially only focused on students as participants, we found that faculty and staff were also interested, often because they were not able to do lab or field-based research.

We determined participants’ coding experience prior to SCIP and coding confidence before and after SCIP. Here, we report only on the participants for which we could pair the preassessment and post-assessment surveys. In 2020, we were able to pair 75 pre- and post-assessments out of 112 pre-assessment surveys. In 2021, we were able to pair 63 pre- and post-assessments out of 128 pre-assessment surveys (Figure 6). 7 paired survey observations were excluded in Summer 2021 data due to the participants taking part in the Image Processing groups that did not learn coding skills. The majority of the participants assessed had no or only some experience with coding in both cohorts: 50.7% and 50.8% had no experience in coding, and 18.7% and 31.7% had less than 1 semester of experience, in 2020 and 2021 respectively (Figure 6).

**Figure 6.**
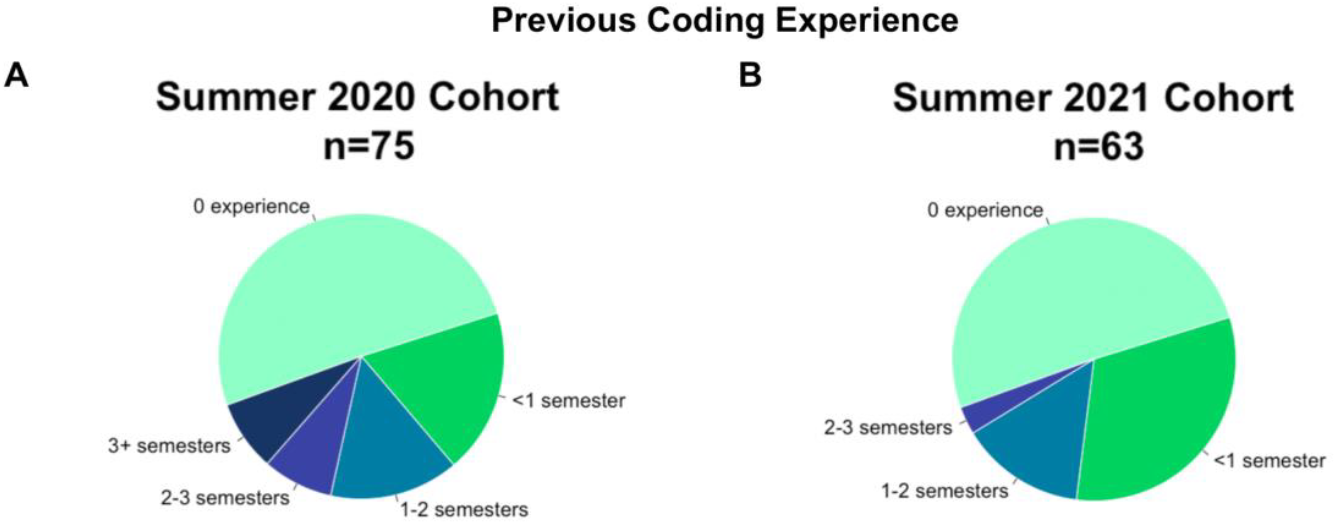
Survey-reported experience levels of participants prior to attending SCIP in 2020 and 2021.

To find out if the SCIP program influenced participants’ coding confidence, we used self-assessment in the pre- and post-assessment surveys to find out how confident participants felt about their coding skills on a Likert rating scale from 1 through 5. Overall, the results showed that participant coding confidence increased (Figure 7).

**Figure 7.**
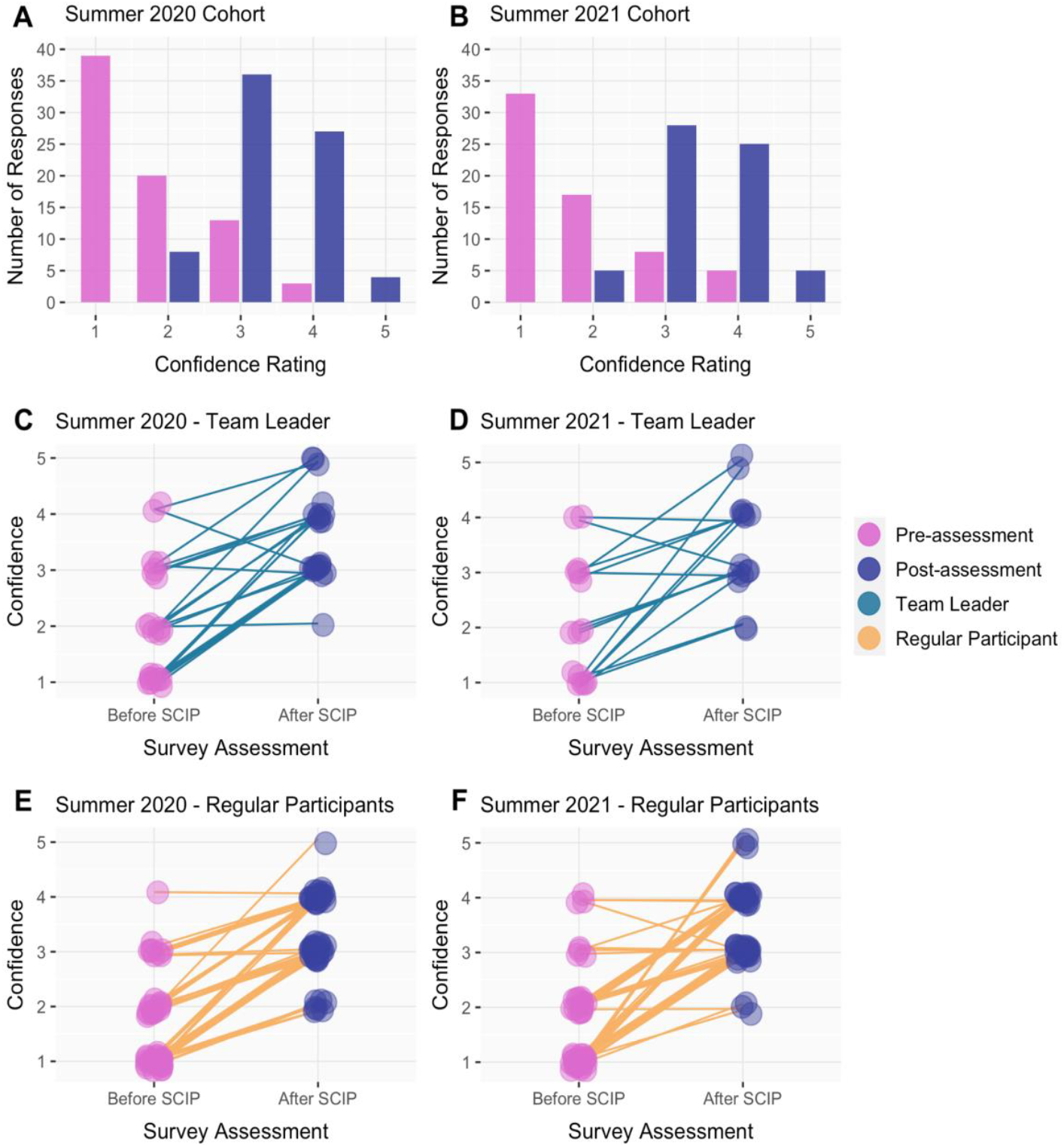
Coding confidence of 75 participants with paired survey responses for pre- and post-assessments in 2020 and 63 participants in 2021. **(A/B)** The number of responses per rating based on the question “Before SCIP/After SCIP, how confident did you feel in your coding skills?” for 2020 and 2021, respectively; **(C/D)** Paired survey responses of participants who identified as team leaders. Each line is an individual participant indicating coding confidence prior to (pink) and after (purple) the completion of SCIP for each year; **(E/F)** Paired survey responses of participants who were regular participants. Each line is an individual participant indicating coding confidence prior to (pink) and after (purple) the completion of SCIP for each year.

For both SCIP Summer 2020 and 2021 the median coding confidence for all participants increased 2 points - from 1 in the pre-assessment to 3 in the post-assessment - with a p-value of 4.32e^-16^ (df=4) and n = 75 for Summer 2020 and a p-value of 3.71e^-14^ (df=4) and n = 63 for Summer 2021. Since the p-values are less than 0.05, we can infer that the increase in coding confidence was significant (Figures 7A, B).

To determine if there was a difference in coding confidence change in team leaders and in regular participants, we analyzed these groups separately. For team leaders, the median coding confidence for both cohorts increased by 1 point - from 2 in the pre-assessment to 3 in the post-assessment - with a p-value of 1.65e^-04^ (df = 4) and n = 23 for Summer 2020 and a p-value of 0.04 (df=4) and n = 15 for Summer 2021. Based on the p-value, we can infer that team leaders had a significant increase in median coding confidence. Next, we analyzed regular participants’ median coding confidence. The median coding confidence for regular participants increased 2 points in both cohorts - from 1 in the pre-assessment to 3 in the post-assessment - with a p-value of 1.07e^-11^ (df=4) and n = 52 for Summer 2020 and a p-value of 1.03e^-12^ (df = 4) with an n = 48 for Summer 2021. Based on the p-value, we can infer that there was a significant increase in median coding confidence in regular participants (Figures 7C-F).

Of the participants who completed a post-program survey, over 75% of individuals were planning or considering taking a coding class in the future. A large majority of our participants had little-to-no experience with coding prior to the start of the program. Of those with little-to-no experience, the majority of participants responded that, after the program, they were considering or planning to take a class with a coding component in order to further their coding experience (Figure 8).

**Figure 8.**
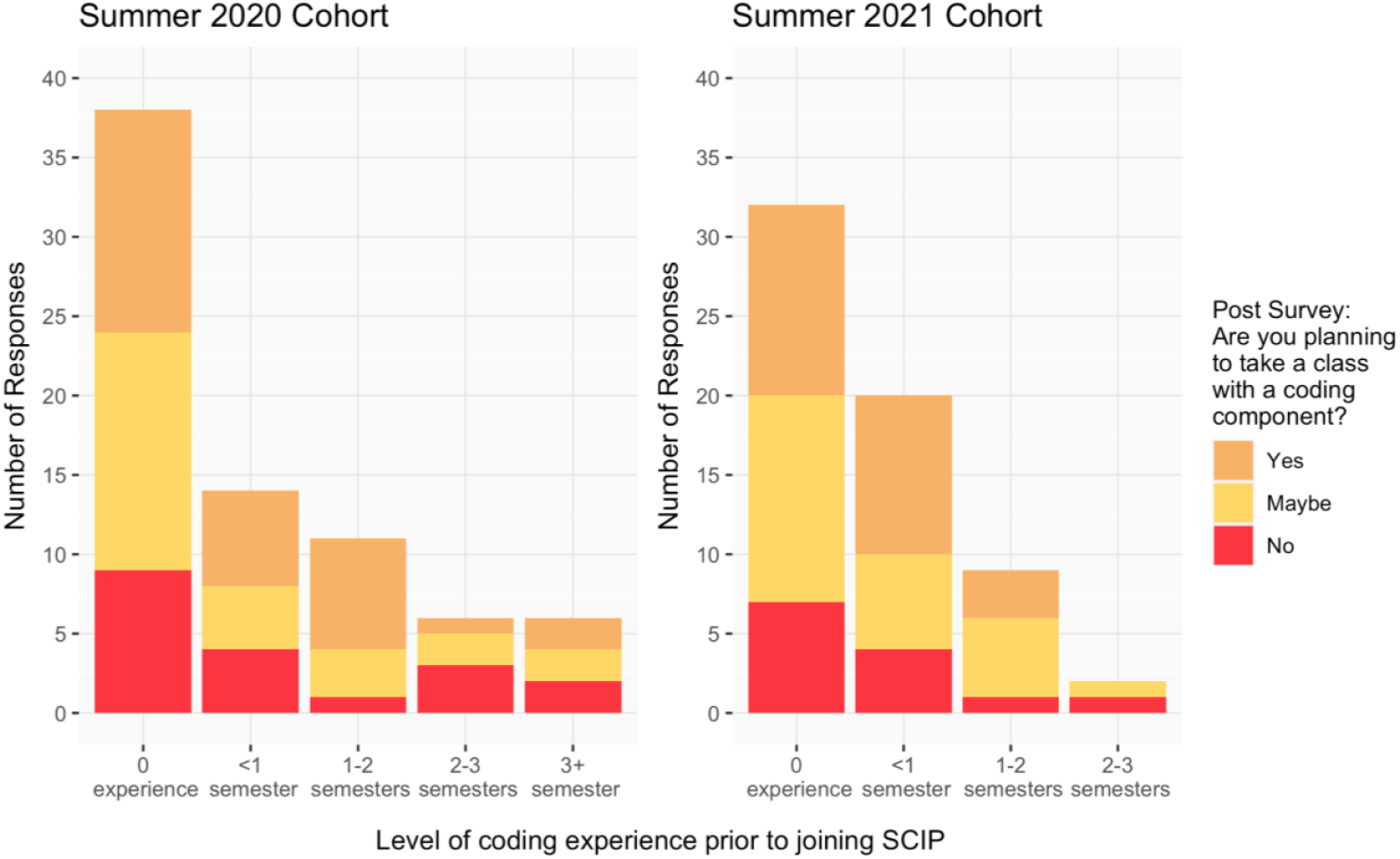
Participants’ survey responses based on the question, “Are you planning to take a class with a coding component?”, regarding their interest in taking a class with a coding component after SCIP from Summer 2020 and Summer 2021.

## CONCLUSION

The impact of the pandemic on our student population included loss of income, loss of family members, and loss of scientific experiences due to remote learning. We developed a virtual program that provided students with opportunities to learn a new skill and acquire confidence in this skill, while also building a community from behind a computer screen. Two faculty and three part-time staff members managed over 130 participants in this virtual environment for both 2020 and 2021. Retention was high over the 6–8-week period and participant feedback was positive. Coding confidence increased significantly, and the participants appreciated the community and support they found in the self-paced program.

Participant comments in the open-ended questions suggested that the community was an important part of the learning environment: *“I liked the fact that my team was open to communicating in and out of zoom. I felt comfortable talking to them and asking them questions”, “I liked that we all had fun doing it, struggling, teaching each other”* and “*It was a safe supportive environment: everyone was really nice. I liked getting to know my team members*”. With more than a year of challenges associated with the pandemic, it has become clear that the challenges of shifting from in-person learning to online learning, including the loss of summer research opportunities, have been far greater than just technical adjustments. Studies worldwide have shown that the lockdown associated with the COVID-19 pandemic has had many impacts on student mental health, including changes in eating behaviors and sleep, and increases in anxiety, depression and suicidal thoughts (Flaudias et al., 2020; Kaparounaki et al., 2020; Son et al., 2020; Wathelet et al., 2020; Yang & Ma, 2020). Given the role that isolation has on mental health in general and its particular impact on students during the pandemic, through isolation from social networks, a lack of interaction and emotional support, as well as physical isolation (Elmer et al., 2020), it will be important to assess the impacts of a program like SCIP on the mental health of students.

Finally, as science becomes more interdisciplinary, coding and data science are clearly taking a more prominent role in biological and biochemistry research. The increase in confidence gained by our participants, many from historically underserved groups, is motivating for us to create more racial diversity in fields not typically diverse, but where these computational skills are particularly important - such as neuroscience, computational biology, evolutionary biology, and ecology. In the future, we aim to offer this program to more students and hope that other campuses will learn from our program. The SCIP method can be used within specific research fields, to draw students from across the country with this free format.

Our future goals are to create new online courses with diverse representation amongst the course instructors to offer as part of future SCIP programs, as a mechanism to increase coding skills amongst our students and to increase diversity in computational fields.

## Supporting information

Supp file 1 Registration

Supp file 2 Survey questions

Supp file 3 Team leader information

SupplementaryFigure1&2_Major_AcademicLevel_SlackActivity

## ACKNOWLEDGEMENTS

This work was supported in part by the NIH MBRS-RISE (#R25-GM059298), NIH MS/PhD Bridge to Doctorate (#R25-GM048972/#T32-GM142515), NIH MARC (#T34-GM008574), NIH URISE (#T34-GM145400), NIH Bridge to Baccalaureate (#5R25-GM050078), NSF-STC CCC (#DBI-1548297), Genentech Foundation Scholarship (#G-7874540), NSF-REU (#DBI-1659175), UCSF-SFSU Cell Design Institute Scholarship and BMS BE-Stem Scholarship programs. We would also like to thank Dr. Senay Yitbarek, Dr. C. Brandon Ogbunu and members of Dr. Pennings’ CoDE Lab for their support. We do not have any conflicts of interest to declare.

