## Supplementary material for "SCIP: a self-paced, community-based summer coding program creates community and increases coding confidence, lessons learned from the pandemic": Supp file 1 Registration

### Join the Science Coding Immersion Program

NOTE: deadline to sign up is May 15, 2021. Once you sign up, we'll let you know whether we have a spot for you. Please keep a look out for an email every week from us to confirm that we have received your email. The program starts on June 1, 2021.

If you are changing responses in your form, please do NOT send in another registration form. If you have previously applied for the program, you are able to edit your submitted form via a link through the response receipt that was emailed to you after you submitted your application. Please email us at if you have any questions regarding this!

----------------------------------------------------------------------------------------------------------------------------------------------------------------------------------------

The Science Coding Immersion Program (SCIP) is run by the Code Lab @SFSU through Dr. Pleuni Pennings, Ryan Fergusson, Olivia Pham, and Rochelle-Jan Reyes and also supported by Dr. Sarah Cohen and Dr. Megumi Fuse. Our goal is to help many students to learn coding skills during the Summer of 2021.

Why a coding program for students in Biology and Biochemistry?

Coding is a useful skill for scientists, but it is hard to learn alone and not every school has nice coding classes for biologists. That's why we want to help you this summer. If you dedicate 9 hours a week to coding (8 hours with your team and 1 hour for a webinar every week), we believe that you can learn a lot of coding skills – even if you are an absolute beginner. Plus, it'll be fun because you're working in a team.

We hope that our program will give you a friendly community and hope for a better future.

**Important things to know about the program**
1. You don't need any prior experience with coding or computer science (but if you have some, you are welcome too).
2. This program is FREE!
3. The program is 9 hours per week for 6 weeks.
4. You'll be learning the language R or Python or Image Processing/ImageJ.
5. This is an immersion program. You will spend 8 hours per week on zoom with your team. There's no homework outside these hours.
6. This is not a class. Therefore, it's not graded nor does it provide any form of educational credit.
7. You will work in a small team (5-6 students) with a team leader.
8. If you are interested to be a team leader, please indicate on this form!

Every team meets for 2 hours every Monday through Thursday, starting from June 1 and ending on July 9, 2021. Participants will also meet 1 hour on Thursdays for the weekly webinars. If you registered for the program before or by April 15, you will receive a secondary link to determine your preferred schedule.

Please fill out this Google form if you are interested in joining the Science Coding Immersion Program!

Best,
Pleuni, Ryan, Rochelle, Olivia (for questions, please)

* Required

1. Email *

2. What is your first name? *

3. Last name? *

4. Preferred name (if different from your first name)

5. How much coding experience do you have? (No experience is needed for this program) *

   *Mark only one oval.*

   - No experience at all
   - I know a few commands here and there (Some experience, <1 semester)
   - I have a handle of the basics (Average experience, 1-2 semesters)
   - I have some experience programming (More than average experience, 2-3 semesters)
   - I'm a pro! (Lots of experience, 3+ semesters)

6. What university/college/school are you attending? (Please put the full university/college/school name, no abbreviations please) *

7. What is your major? Add concentration, if applicable. *

8. What is your current education level? *

*Mark only one oval.*

- Undergraduate student
- Just graduated, not in a program yet
- Post-bacc student
- Master's student
- PhD student
- Post-doc
- Staff / faculty / lecturer
- Other: ____________

9. Are you currently in one of these programs? *

*Mark only one oval.*

- Promoting Inclusivity in Computing (PINC) - for SFSU undergraduate students
- Graduate Opportunities to Learn Data Science (GOLD) - for SFSU graduate students
- I am not in either

10. Which race / ethnicity do you identify with? Choose all that apply. *

*Check all that apply.*

- [ ] American Indian or Alaska Native
- [ ] Asian
- [ ] Black or African American

- [ ] Native Hawaiian or Pacific Islander
- [ ] Hispanic, Latino, or Spanish Origin
- [ ] Not Hispanic, Latino, or Spanish Origin
- [ ] White
- [ ] Prefer not to say

#### Info for the Program

Every team meets 2 hours every Monday through Thursday, starting from June 1 and ending on July 13, 2021. Participants will also meet 1 hour on Thursdays for the weekly webinars. If you register for the program before or by April 15, you will receive a secondary link with your confirmation to determine your scheduling availability!

11. Have you participated in SCIP (Summer 2020 or Fall 2020) before? (If so, make sure to not choose the same coding language this year.) *

*Mark only one oval.*

- ( ) Yes
- ( ) No

12. In the SCIP program you can learn R or Python (these are coding languages that are used often in biology and chemistry). Which would you like to learn? *

Note: if you have taken CSC 306 (an SFSU course designed for biology and chemistry undergraduates to learn programming via Python), we recommend that taking the R course unless you are interested in a refresher for basics of Python. This is because SCIP is designed for those who have no prior experience and would like to learn the basics for each language.

*Mark only one oval.*

- ( ) I'd like to do the Beginners R course

I'd like to do the Beginners Python course

I don't have a preference

13. Would you be interested to be part of a pilot and do a beginners image processing course (using image J) [limited spots available]? *

*Mark only one oval.*

Yes

No

14. You'll be working in a team with 4-6 students. Would you be willing to be a team leader? This means that you facilitate team meetings on Zoom. If you have experience managing a team, for example in your job, sorority, church or dorm, you can be a team leader. *

Please note that you do NOT need to have coding experience prior to the program to be a team leader!

*Mark only one oval.*

Yes

No

15. If you are interested to be a team leader, please describe your experience working with people and/or facilitating meetings. (1-3 sentences) *

Time Availability for Team Meetings & Webinars

These questions will delve into your preferences for times to determine what group would best suit your choices. Please indicate ALL of the meetings times you are available for, and one will be selected.

16. TEAM MEETINGS: Are you available to meet M, T, W, Th 9-11am? *

*Mark only one oval.*

- Yes
- No

17. TEAM MEETINGS: Are you available to meet M, T, W, Th 12-2pm? *

*Mark only one oval.*

- Yes
- No

18. TEAM MEETINGS: Are you available to meet M, T, W, Th 4-6pm? *

*Mark only one oval.*

- Yes
- No

19. TEAM MEETINGS: Are you available to meet M, T, W, Th 7-9pm? *

*Mark only one oval.*

- Yes
- No

20. WEBINARS: Are you available to attend a webinar on Th 11am-12pm? *

*Mark only one oval.*

- Yes

[illegible]

No

21. WEBINARS: Are you available to attend a webinar on Th 2-3pm? *

*Mark only one oval.*

Yes

No

22. WEBINARS: Are you available to attend a webinar on Th 3-4pm? *

*Mark only one oval.*

Yes

No

#### Additional Info

23. What kind of operating system would you use for the program? *

*Check all that apply.*

- [ ] Mac (desktop or laptop)
- [ ] PC (desktop or laptop)
- [ ] Tablet (Apple or Android)
- [ ] Phone (Apple or Android)

24. Do you have reliable WiFi or mobile hotspot for attending Zoom meetings? *

*Mark only one oval.*

Yes

[illegible]

No

Other: ______________________

25. How did you find out about this program? *

*Mark only one oval.*

- Social Media (Twitter, Instagram, Facebook, LinkedIn)
- PINC Website
- Heard from the Biology and/or Biochemistry department at SFSU
- Through the CCC Program
- From the SEO office
- Announcement in a class
- Other: ______________________

26. Do you have any questions for us?

This content is neither created nor endorsed by Google.

Google Forms
