## Supplementary material for "SCIP: a self-paced, community-based summer coding program creates community and increases coding confidence, lessons learned from the pandemic": Supp file 2 Survey questions

### Science Coding Immersion Program – List of Survey Questions

*Please note: all questions were used in both 2020 and 2021, unless stated in parentheses prior to the beginning of the survey question.*

**Mid-Program Survey (2020) & Pre-Program Survey (2021)**

Prior Coding Experience

1. Prior to joining SCIP, what was your level of coding experience?
2. (2021) If you have previous coding experience, what language did you use?
3. (2021) If you have prior coding experience, where/how did you learn to code?
4. (2020) Prior to SCIP, did you take CSC 306 (Intro to computing class which is part of the PINC program)?
5. Prior to joining SCIP, how confident did you feel in your coding skills?
6. (2021) What do you want to use your programming skills for?

Your SCIP Experience

7. (2020) How many weeks are you into SCIP now?
8. Are you a SCIP team leader?
9. Only for team leaders: Prior to beginning the SCIP Zoom meetings with your team, how confident did you feel about being a SCIP team leader?
10. (2020) If you are not the team leader, who is your team leader?
11. (2020) How effective is/was SCIP Zoom meetings with your team?

Personal Demographics

1. (2020) Are you an SFSU student?
    a. *Changed to:* (2021) What College or University do you currently attend?
2. (2020) If you are an SFSU student, are you SEO supported?
3. (2020) Are you an undergraduate or graduate student?
    a. *Changed to:* (2021) What is your current academia level?
4. (2020) Are you a biology or (bio)chem major?
    a. *Changed to:* (2021) What is your major/degree?
5. (2021) What is your degree concentration?
6. Are you currently doing research or participate in a research lab?
7. What race/ethnicity do you identify with (choose all that apply)?
8. (2021) If you would like to specify race/ethnicity, please list it below.
9. To which gender identity do you most identify?

Anonymous Identification

10. What is your address number (number only)?
11. What is the first letter of your middle name (or another memorable letter for you)?
12. What are the last four digits of your phone number?

---

**Post-Program Survey (2020 & 2021)**

(2021) Agreement

1. (2021) By completing the survey and choosing the “I accept” button below, you are allowing the SCIP Admin Team to utilize this information for review and data analysis purposes. However, your responses are and will be 100% anonymous and cannot be – in any way, shape, or form – tracked to you.

Coding Experience

1. (2021) What course did you participate in?
2. After completing SCIP, how confident do you feel in your coding skills (or image analysis skills) overall?
3. How comfortable do you feel in your coding abilities to create a graph from data?
4. How confident do you feel in your coding skills to mutate/change data within R or Python?
5. Are you planning to take a class with a coding component?
    a. *Changed to:* (2021) In the upcoming semesters, are you planning to take a class with a coding component?
6. (2020) Would you be interested in participating in SCIP next summer?
7. (2021) Do you plan for coding to be part of your future career?

Team & Community Experience

8. Are you a SCIP team leader?
9. TEAM LEADERS ONLY: How confident did you feel about being a team leader towards the end of the program?
10. If you are not the team leader, who is your team leader?
11. How effective is/was SCIP Zoom meetings with your team?
12. Did you feel supported by your team and your team leader through the program?
13. Did you feel comfortable asking questions during your Zoom meetings?
14. How effective was having Zoom meetings - 2 hours/day for 4 days/week – for your learning-to-code experience?
15. SHORT ANSWER: What did you like about the Zoom meetings with your team?

Udacity Course & Learning

16. (2020) What Udacity Course did you take?
17. Approximately, how far did you reach in the Udacity course?
18. Did you use extra resources in order to learn Python or R, aside from the Udacity course resources and SCIP-provided readings?
19. Would you recommend the Udacity course to others?
20. SHORT ANSWER: If you do not recommend the Udacity course, please explain why and if there are any other courses you may recommend.
21. (2021) R TEAM MEMBERS ONLY: What is one thing you liked about the R videos by Ryan F?
22. (2021) R TEAM MEMBERS ONLY: What is one thing we could improve about the R videos by Ryan F?
23. (2021) R TEAM MEMBERS ONLY: What is one thing you liked about the coding projects?

Overall SCIP Experience

24. (2021) What was/were the reason(s) you chose to join SCIP?
25. (2021) Were there other reasons you chose to join SCIP not listed above?
26. (2020) How many weeks did you stay in the program?
27. SHORT ANSWER: If you stopped attending the program at any point, please explain why you stopped attending.
28. How useful was Slack for you and/or your team?
29. How useful were the SCIP webinars?
30. Please explain your answer from the previous question regarding the SCIP webinars.
31. Did you feel supported by the SCIP Administrative Team?
32. Would you recommend SCIP to others?
33. SHORT ANSWER: What is one thing you liked about the program?
34. SHORT ANSWER: What is one improvement you would like to be implemented for this program in the future?
35. SHORT ANSWER: Please let us know any other comments or suggestions you may have about the program!

Anonymous Identification

36. What is your address number (number only)?
37. (2020) What is the first letter of your middle name (or another memorable letter for you)?
38. What are the last four digits of your phone number?
