## Supplementary material for "SCIP: a self-paced, community-based summer coding program creates community and increases coding confidence, lessons learned from the pandemic": Supp file 3 Team leader information

### Science Coding Immersion Program

Team Leaders - Summer 2021 Cohort

Thank you again for choosing to become a team leader! You will be a crucial source of support not only for the program but also for your team.

In this document, we will provide you all the information you will need in order to start your team meetings and create a supportive environment open for discussions!

#### Roles & Responsibilities

##### 1. Creating a Zoom Meeting Link

Your first responsibility in the program is to create a Zoom meeting link for your team. This will be your team's meeting location for the entirety of the program. In the event of webinars, a meeting link will be provided to everyone.

To create a Zoom meeting link, first you must download zoom: use this https://zoom.us/download → click "Zoom Client for Meetings". After downloading and opening Zoom, sign in with your primary email address. After signing in, click the drop down menu of "New Meeting" and click "Copy Invitation". This creates a Zoom meeting link.

##### 2. Lead the Team Meetings

Your primary role as team leader during meetings is *not* to instruct your team on the lesson itself, but rather to guide your team through discussion. In this way, your role is similar to that of a near-peer mentor. You are learning the course *with* your team, rather than teaching them. As team leader, your two primary responsibilities are to ensure that everyone either understands the lesson/video presented, or identify the confusions about the lesson/video. You do this by facilitating discussion with your team.

> *Suggestion: This could be in the form of 5-minute check-ins during your team meeting to ask where everyone is at and if anyone is having difficulty understanding anything.*

Your secondary role as team leader is that of time manager for meeting activities. Your team meeting will often involve activities such as weekly readings (pg. 6), or reviewing resources shared by the Admin Team. We provide suggested formats of how to structure your meeting with these activities (pg 4).

Leading team meetings has other responsibilities as well. The responsibilities include, but are not limited to:

- **Ensuring meetings stay on time:** These meetings often come at the expense of someone's schedule. It's important to respect everyone's time and everyone's effort to be punctual.
- **Recording absences:** We won't penalize a small number of absences, but it's important for the Admin Team to know who's coming every week and who hardly shows up.
- **Creating breakout rooms or polls:** A situation may occur where just one student is stuck but doesn't want to slow the rest of the team down. Often the team is comfortable explaining the material to the other member, but in the event that they would like to move forward, you can create a breakout room so someone can explain the problem in a one-on-one setting.
- **Gathering questions:** In the event that the whole team encounters a misunderstanding, you may gather questions to post onto the **#question-forum** Slack channel. This gives the Admin Team an opportunity to address your question, or possibly give a demonstration during one of your meetings.

#### 3. Ensure Survey & Form Completion

From time to time, the SCIP Admin Team will be requesting that each SCIP participant complete a form or survey. As a team leader, you are responsible for reminding your team to complete these documents and ensure that they are completed. The most important documents will be **the Honor Code Agreement & Audio/Visual Release Form, the Pre-Program Survey, and the Post-Program Survey**. *These reminders can be a simple message in your Slack team channel or an announcement during your meeting.*

**Summer 2021 SCIP Honor Code Agreement & Photo Consent Form link**:
https://forms.gle/Rn1p23iiKMaA4VSA7

**Summer 2021 SCIP Pre-Program Survey link**: https://forms.gle/3H739WRyh8VDuEEp6

#### 4. Learn R, Python, or ImageJ!

Last but not least, one of your duties is to learn R, Python, or ImageJ! Although team leaders will have some extra responsibilities, you are also a participant of the program learning how to code. We want to ensure that you are also learning R, Python, or ImageJ during SCIP. If you need more support aside from your team, please reach out to any member of the Admin Team.

### The First Meeting

#### 1. Sending the Zoom Link Invitations

In the first week of the program, your main tasks are to create a Zoom meeting link and guide your team through course familiarization. Before your team can meet, the Zoom link invitations must be sent.

**Sample email to invite your team:**

> Dear [NAMES OF TEAM MEMBERS],
>
> I am your team leader for the Science Coding Immersion Program this Summer 2021. Below, I put the information for our first Zoom meeting:
> [INSERT ZOOM MEETING INFO HERE]
>
> It will take place on Monday - Thursday at [ADD TIME]. I am excited to be working with you the next 6 weeks!
>
> Best,
>
> [YOUR NAME]

#### 2. Leading Your First Meeting

Now you're ready to start leading meetings for SCIP! Following the suggested guideline for Day 1 will guarantee that you're prepared to manage the time for the first meeting. Given that this will be the beginning of the program, the format of the first meeting will vary significantly from the format of subsequent meetings.

| Day 1 | |
|---|---|
| **Introductions**<br>15 min | Follow the following prompts:<br>● What is your name?<br>● What is your major? What is your career goal?<br>● Why are you interested in learning coding skills?<br>● Favorite take-out place? Favorite artist on Spotify at the moment? |
| **Form Completion**<br>10 min | Check-in to see if everyone has completed the following docs and give the team some time to complete this if they haven't.<br>● **Summer 2021 SCIP Honor Code Agreement & Photo Consent Form link**: https://forms.gle/Rn1p23iiKMaA4VSA7<br>● **Summer 2021 SCIP Pre-Program Survey link**: https://forms.gle/3H739WRyh8VDuEEp6 |

| | |
|---|---|
| **Join Slack Workspace**<br>5 min | • Check in to see if everyone joined the SCIP Summer 2021 Slack workspace<br> ○ **Invitation for the Slack workspace:** https://join.slack.com/t/sfsu-scip-su21/shared_invite/zt-qhlllv91-vBy_ZaXeKlV9~IZSM4zRDg<br> ○ Refer team members to this video if they're unfamiliar with Slack: https://vimeo.com/427096842/ba37af3e53<br>• Make sure everyone is familiar with all Slack channels. To review the Slack channels, please review the "2021 Program Information". |
| **Orientation Video**<br>5 min | • This video will introduce the SCIP Admin Team and welcome you into this year's Science Coding Immersion Program!<br> ○ https://vimeo.com/554549535/4c83d53a55 |
| **Join Online Coding Class**<br>15 min | Dependent on what programming language your team is learning, each of you will need to sign up and create an account to take the online coding class:<br>• **R**: https://www.udacity.com/course/data-analysis-with-r--ud651<br>• **Python**: https://www.udacity.com/course/introduction-to-python--ud1110<br>• **ImageJ**: https://www.edx.org/course/image-processing-and-analysis-for-life-scientists<br><br>Sign up for the course and make sure everyone has signed up as well. Please refer to the **course guide** to see suggestions for when to complete the modules.<br><br>*Suggestion: You can initially all watch the videos from the team leader's computer and do the quizzes together. After your meeting, send us a message to let us know what's not working.* |
| **Working Time**<br>45 min | Quiet working time to go through the first few videos. Depending on your team's learning style, teams can utilize this time to go through the course together (i.e. team leader shares their screen and audio) or separately but muted (i.e. each participant watches the videos on their own in the meeting).<br><br>*Suggestion: Team members can mute when working on the course, but can unmute and ask for help if they have any questions.* |
| **Discussion**<br>15 min | • Discuss what you learned from the first videos and quizzes.<br>• Make a list of 2 things you learned and 1 thing you'd like to learn. |
| **Questions**<br>10 min | Collect any questions and post them on the **#question-forum** Slack channel. |

### Regular Meetings

#### 1. Meeting Formats

Decide whether your team prefers a 1-hour block of working time (Option 1) or two 30-minute blocks of working time (Option 2) with a break in between. If there is another format that works best for your team that covers the following daily activities, then you are free to use it.

<table>
<tr><th colspan="2">Option 1</th><th colspan="2">Option 2</th></tr>
<tr><td>15 min</td><td>Check ins: Discuss the previous open questions & everyone’s progress with course materials.</td><td>15 min</td><td>Check ins: Discuss the previous open questions & everyone’s progress with course materials.</td></tr>
<tr><td>15 min</td><td>Weekly Readings: Suggested reading (see list below) and discussion (in Break-Out Rooms or full group)</td><td rowspan="2">30 min</td><td rowspan="2">Quiet Working Time-1: Students can unmute for brief questions.</td></tr>
<tr><td rowspan="3">60 min</td><td rowspan="3">Quiet Working Time: Students can unmute for brief questions.</td></tr>
<tr><td>15 min</td><td>Weekly Reading: Suggested reading (pg 4.) and discussions in Break-out Rooms or all group</td></tr>
<tr><td>30 min</td><td>Quiet Working Time-2: Students can unmute for brief question</td></tr>
<tr><td>15 min</td><td>Questions: Identify confusions and discuss. If something remains unclear, post questions to #questions Slack channel. The SCIP team loves questions!</td><td>15 min</td><td>Questions: Identify confusions and discuss. If something remains unclear, post questions to #questions Slack channel. The SCIP team loves questions!</td></tr>
<tr><td>15 min</td><td>Reflective Journaling & Closing: Here is a short video on journaling in SCIP<br><br>https://www.powtoon.com/s/coWmq2yiKJC/1/m</td><td>15 min</td><td>Reflective Journaling & Closing: Here is a short video on journaling in SCIP<br><br>https://www.powtoon.com/s/coWmq2yiKJC/1/m</td></tr>
</table>

#### 2. Reflective Journaling

Reflective journaling has been shown to help with learning and to stay motivated. We highly recommend doing it at the end of each meeting. The team members should write in a notebook or their computer about their learning experience that day. Reflections help everyone get the most out of the lessons. It provides each team member with time to consider what they are learning and why. Each day, you can use one or more of these prompts (or you can share them all and let the team members decide too) :

- **Today I learned about…**
- **I am still confused about…**
- **Today I am proud of / Today I am thankful for…**
- *Suggestion: we recommend leaving at least 8 minutes for team members to reflect quietly*

#### Weekly Readings

Each week, you will take one of the 15 minute time blocks in your meeting to read an article or page. Topics will vary, but below are readings you will use for these time blocks. Below is the guide for weekly readings.

| | | | |
|---|---|---|---|
| **Week 2 Readings** | | | |
| Scientist Spotlight: Berenice Chavez Rojas (Link) | COVID-19 project from SCIP 2020 by Johnny Duong (Link) | Programming tips for beginners (Link) | Self-care for the scientist (Link) |
| **Week 3 Readings** | | | |
| Scientist Spotlight: Jazlyn Mooney, PhD student UCLA (Link) | Best Practices for Scientific Computing (Link) | Wu and Watterson's Theta - Research done with coding (Link) | Four tips to ward off imposter syndrome (Link) |
| **Week 4 Readings** | | | |
| Scientist Spotlight: Meet Dr. Yee Mey Seah (Link) | Ten Rules for Coding (Link) | Understanding our eugenic past to take steps towards scientific accountability (Link) | Math in biology by Dr Scott Edwards (Harvard professor) (Link) |
| **Week 5 Readings** | | | |
| Scientist Spotlight: Dr. Ryan Hernandez (Link, Video) | R vs Python (Link) | GirlsWhoCode Chapter 2 (Link) | 1 woman's mission to empower Ethiopia's youth thru tech (Link) |
| **Week 6 Readings** | | | |
| Scientist Spotlight: Meet Dr. Sabah Ul-Hasan (Link) | Studying Computer Science (Link) | "What is bioinformatics?" By Maria Nattestad (Link) | Ways to look after yourself and others in 2021 (Link) |
| **Programming Language-Specific Readings** | | | |
| Why should a biologist learn R? (Link) | 5 Reasons why you should learn Python (Link) | What Language to Start With - Python (Link) | Rosana Callejas about Learning R (Link) |

| Extra Readings | | | |
|---|---|---|---|
| Visualizing Amounts (Link) | A gentle intro to machine learning for bio and chem students (Link) | Scientist Spotlight: Graham Larue (Link) | Visualizing Geospatial Data (Link) |
| Scientist Spotlight: Meet Francisca Catalan, SFSU PINC alum & research associate at UCSF (Link) | 10 Rules of studying every software developer should follow (Link) | The math of Ferguson (Article; RMD File) | Scientist Spotlight: Meet Simone Webb, Bioinformatics and Immunology PhD student (Link) |

| Extra Readings (Image Analysis) WEBINAR June 17, 2021 | |
|---|---|
| The scientific paper applying our image analysis tools and new analyses to a very large 3D image dataset (Link) | The methods paper describing our Image Processing tool/methodology crucial to being able to do the analysis in the scientific paper (Link) |
