## SupplementaryFigure1&2_Major_AcademicLevel_SlackActivity for "SCIP: a self-paced, community-based summer coding program creates community and increases coding confidence, lessons learned from the pandemic"

### Supplemental Tables and Figures

**Table S1**. List of programming languages or softwares we offered as well as the source of these classes.

| Programming Language/Software | Course | URL |
|---|---|---|
| R | Udacity Exploratory Data Analysis with R | https://www.udacity.com/course/data-analysis-with-r--ud651 |
| Python | Udacity Introduction to Python | https://www.udacity.com/course/introduction-to-python--ud1110 |
| ImageJ | edX Image Processing and Analysis for Life Scientists | https://www.edx.org/course/image-processing-and-analysis-for-life-scientists |

**Table S2**. Suggested Zoom meeting format. This table outlines a suggested format that teams could follow during their meeting time. Each team had 4 such meetings each week.

| | **Zoom Meeting Format** |
|---|---|
| **15 mins** | Check ins (make sure everyone talks) |
| **15 mins** | Suggested reading and discussion |
| **1 hour** | Quiet Working Time / Shut Up and Code (this could also two sessions with a break in between) |
| **15 minutes** | Collect questions to answer as a team or post questions and or screenshot of code in “Question Forum” on Slack |
| **15 minutes** | Reflection and journaling |

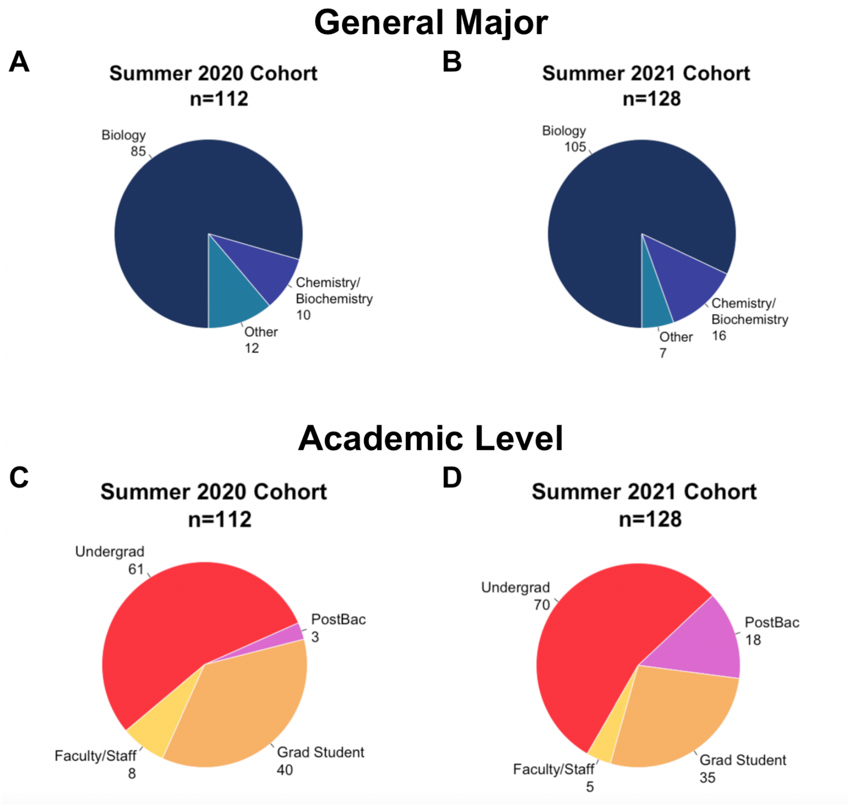


**Figure S1.** Distribution of participants by academic level and major/Department. (A) Respondent-identified majors of SCIP Summer 2020 participants and (B) of SCIP Summer 2021 participants. In Summer 2021, specific majors were asked but general majors were charted. (C) Number of undergraduate, graduate, and post-bacc students as well as faculty/staff that participated in SCIP Summer 2020 and (D) in SCIP Summer 2021.

**Table S3.** Participant Data and Estimates of Participant Retention from Student Communication regarding withdrawal from the program, Survey Responses, and presence in Team Photos. Bolded are the data utilized to calculate the total participation/retention range.

| | **Summer 2020** | **Summer 2021** |
|---|---|---|
| **Starting Number of Participants** | 147 | 138 |
| *Student Communication* | | |
| Notification of Withdrawal | 6 (4.1%) | 11 (8.0%) |
| **Estimated Number of Participants Retained** | **141 (95.9%)** | **127 (92.0%)** |
| *Survey Responses* | | |
| Pre-Survey (% of starting participants) | 112 (76.2%) | 128 (92.8%) |
| **Post-Survey (% of pre-survey respondents)** | **92 (62.6%)** | **76 (55.1%)** |
| *Team Photos* | | |
| **Individuals Pictured in Team Photo (% of starting participants)** | **80 (54.4%)** | **112 (81.2%)** |
| ***Total Participation/Retention Range*** | **80-141 (53.7-95.9%)** | **76-127 (55.1-92.0%)** |

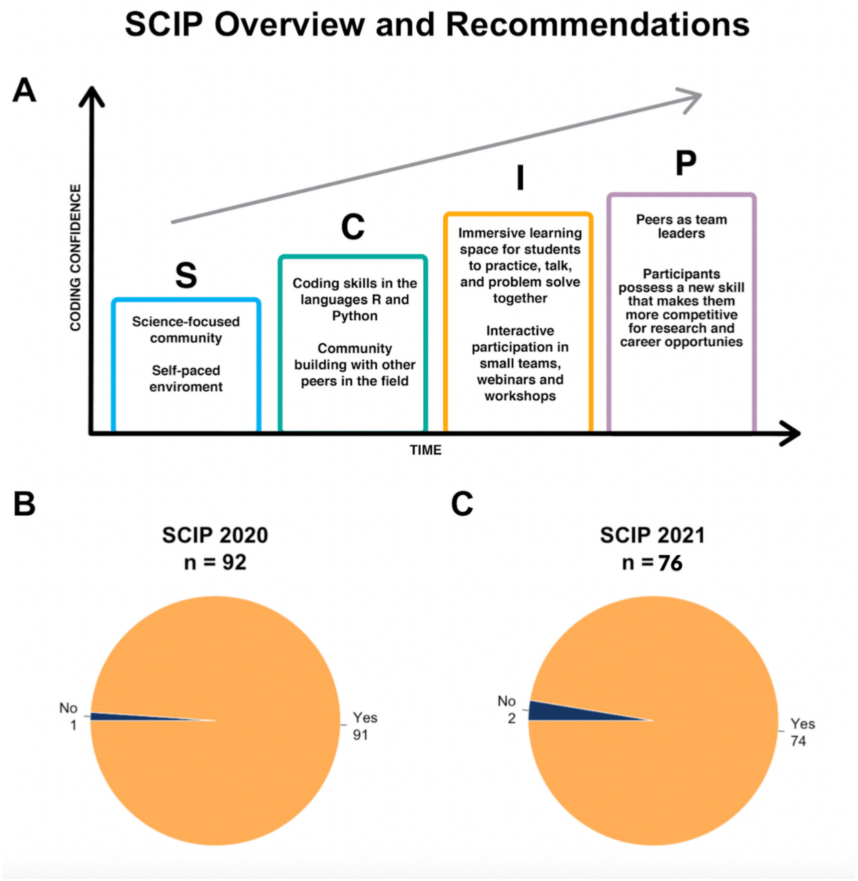


**Figure S2**. (A) Overall plan for design of the Science Coding Immersion Program (SCIP). (B/C) In the post-program assessment, SCIP participants from Summers 2020 and 2021 were asked whether they would recommend SCIP to others. In 2020, 91 out of 92 who answered the question responded that "Yes", they would recommend SCIP, while in 2021, 74 out of 76 answered "Yes".

**Previous Coding Experience**

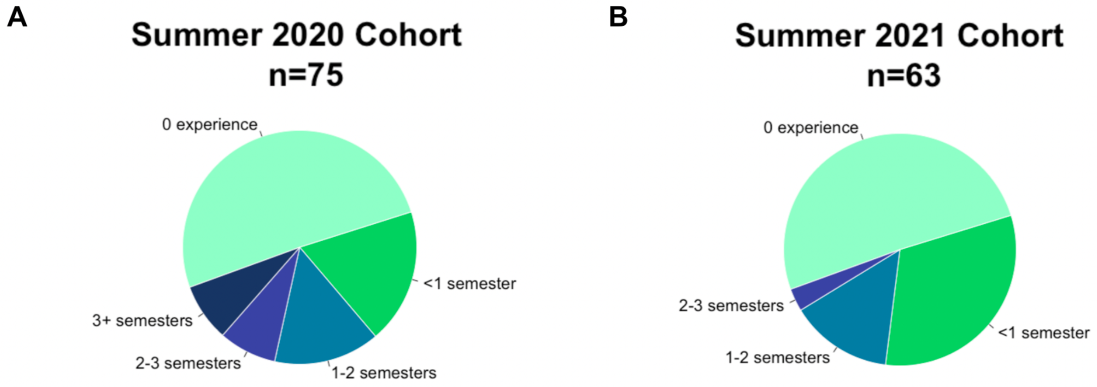


**Figure S3**. Survey-reported experience levels of participants prior to attending SCIP in 2020 and 2021.

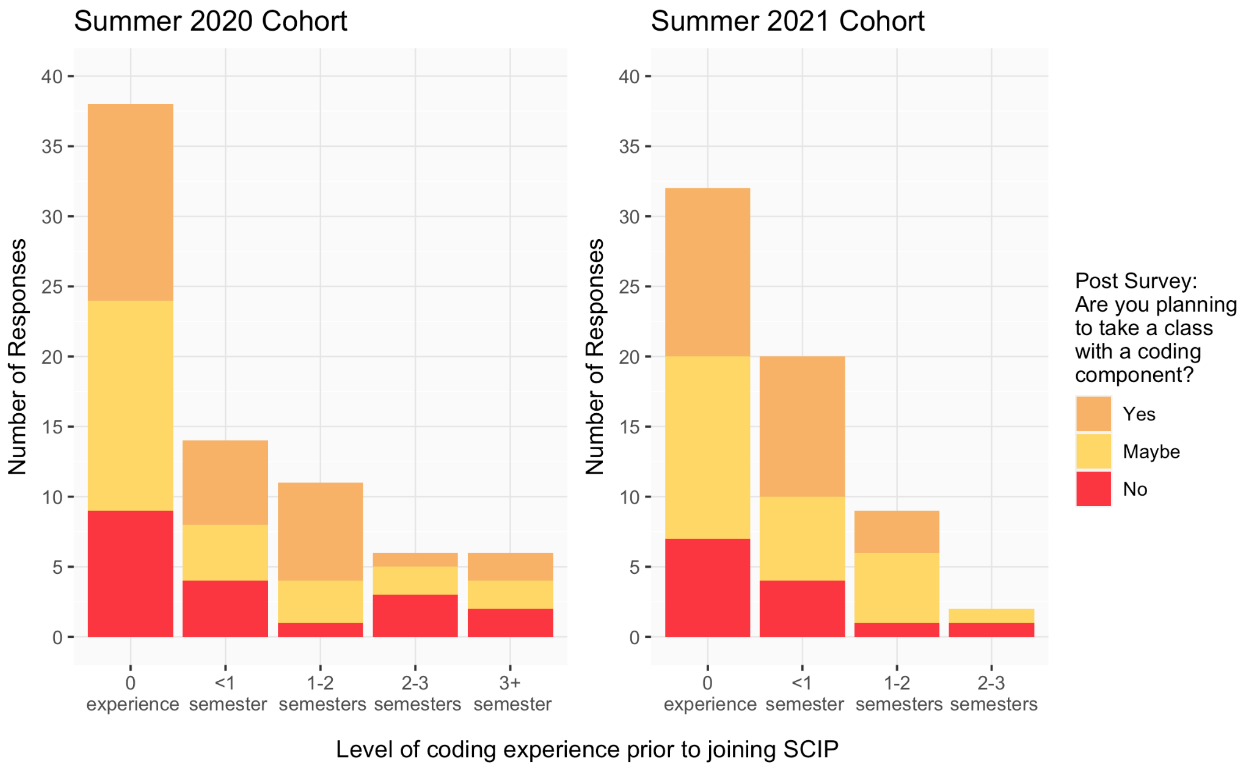


**Figure S4.** Participants' survey responses based on the question, "Are you planning to take a class with a coding component?", regarding their interest in taking a class with a coding component after SCIP from Summer 2020 and Summer 2021.
